# A novel method for investigating fish social behaviour in the field

**DOI:** 10.1101/2022.08.23.504959

**Authors:** Molly A. Clark, Christos C. Ioannou

## Abstract

Field studies of social behaviour are challenging due to having to observe multiple individuals simultaneously. In aquatic environments, these challenges are often amplified by environmental conditions such as habitat complexity, turbidity or darkness, which are often the conditions of interest in the context of studying anthropogenic impacts. Many fish species rely on social interactions for at least part of their life cycle, and while these interactions are known from laboratory studies to be impacted by environmental factors, approaches to quantify social behaviour under natural conditions are limited. Here, we present a novel method whereby multiple funnel traps are deployed simultaneously within a pond to quantify the aggregation and assortment of wild fish populations, using three-spine sticklebacks (*Gasterosteus aculeatus*) as a model species. The number of sticklebacks caught decreased in lower temperatures and through the seasons, from May to November. There was some evidence for decreases in fish numbers with reduced light intensity and higher dissolved oxygen levels. No effect of turbidity was found. The only evidence for changes in aggregation was related to the proportion of breeding males caught, where sticklebacks were less aggregated during the breeding season. These trends are expected based on previous work and knowledge about stickleback biology, validating the method. However, in contrast to previous studies, we found little evidence for phenotypic assortment. This study establishes a new cost-effective technique for investigating the social behaviour of wild fish, with some important benefits over existing techniques.

## Introduction

Social behaviour is essential for the reproduction and survival of many fish species, in at least part of their life cycle. Being in a group provides hydrodynamic (Svendsen *et al*., 2003; Killen *et al*., 2012; Marras *et al*., 2015), reproductive (Taborsky, 2001), foraging (Ranta and Kaitala, 1991) and anti-predator benefits (Ioannou, 2021), as well as costs (Grand and Dill, 1999; Krause and Ruxton, 2002). These costs and benefits associated with shoaling can change depending on environmental conditions, altering group dynamics through adaptive changes in individuals’ behaviour (Pitcher and Parrish, 1993). Environmental conditions can also interfere with and constrain the inter-individual interactions that allows for group formation and maintenance (Chamberlain and Ioannou, 2019; Ginnaw *et al*., 2020). Because of widespread environmental change due to human activity, it is increasingly important to study the effect of environmental factors on social behaviour so we can understand changing dynamics in wild populations, especially in freshwater (Ormerod *et al*., 2010).

Current methods for studying fish social behaviour in the field mostly depend on cameras, or use seine nets to capture whole shoals (e.g. Sarà *et al*., 2007; Croft *et al*., 2009; Burford *et al*., 2022). These methods are often restricted by environmental conditions, particularly those which reduce water clarity, such as turbidity and darkness. Higher light intensity can also produce problems for image and video quality as a result of ‘sunflicker’, increasing visual noise (Gracias *et al*., 2009), and more complex habitats with dense vegetation can add barriers to visibility and create more general access problems. These factors also reduce the ability to analyse videos through tracking software, which generally require a high resolution and contrast between subject and background (Dell *et al*., 2014). Techniques that overcome some of these environmental limitations are often expensive and come with their own constraints. Thermal or sonar imaging allow recording in the field when there is poor water clarity, however have limited spatial and temporal resolutions (Hughey *et al*., 2018; Rodriguez-Pinto *et al*., 2020). Additionally, these technologies present challenges for ID of species, where it is hard to decipher the species being imaged. Alternatively, tracking technologies such as GPS or PIT tags overcome the problems of view, access, and spatial constraints, but require every individual fish to be caught and tagged (Hughey *et al*., 2018). Measuring group behaviour ideally requires data to be collected from a large proportion of the population to avoid missing social interactions, thus these methods involve extensive handling and equipment to reach this goal. These tags can also be invasive with implication for welfare, and it is unclear to what extent they affect the natural behaviours of tagged individuals.

Freshwater ecosystems face multiple threats from environmental change, such as eutrophication and increasing temperatures, and the combined effects of these and other stressors such as overfishing (Carpenter *et al*., 2011; Angeler, 2014; Orr *et al*., 2020). Driven by anthropogenic change, such as urbanisation and agriculture, increased runoff of sediments and pollutants can generate eutrophication and turbidity levels in water that far exceed natural fluctuations (Davies-Colley and Smith, 2001). Elevated turbidity restricts the visual environment for aquatic animals and visibility often deteriorates rapidly, preventing species from being able to adapt to the new conditions (Chamberlain and Ioannou, 2019; Davies-Colley and Smith, 2001). Deforestation can also reduce habitat complexity, leading to fewer foraging opportunities and refuges, and can result in less diverse fish assemblages (Bojsen and Barriga, 2002; Zeni *et al*., 2019). The subsequent reductions in canopy cover from deforestation can also lead to increases in light intensity over streams, creating more noise in the visual environment, e.g. from caustics, and contribute to warmer temperatures (Matchette *et al*., 2018, 2020; Ilha *et al*., 2018). With continued pressure on freshwater environments it is increasingly important to understand how these and other stressors are affecting fish communities and their social behaviour. However, restrictions from environmental conditions on field techniques are often the same environmental conditions that we want to study, such as water turbidity, and novel methods are required to explore their impact on fish social behaviour.

Empirical studies have demonstrated that changes in environmental conditions can impact social interactions in fish. Through masking, environmental stressors can disrupt information transfer between individuals, and thus their ability to maintain coordinated shoals (McNett *et al*., 2010). For example, turbidity and low light intensity restricts the use of visual information among shoal members, leading to reduced group cohesion, coordination, and collective decision making (Pitcher and Turner, 1986; Ryer and Olla, 1998; Ohata *et al*., 2014; Chamberlain and Ioannou, 2019; Ginnaw *et al*., 2020). Consequently, individuals reduce their foraging efficiency and lose the anti-predator benefits provided by shoaling (Chamerlain and Ioannou, 2019). Stressors can also shift focus away from group behaviour via distraction (Chan *et al*., 2010). For example, although there is limited evidence that acoustic cues or signals are used in fish shoaling (Ioannou *et al*., 2011), anthropogenic noise pollution has been shown to shift the attention of individuals, and in turn reduce the coordination and cohesion of shoals (Sarà *et al*., 2007; Herbert-Read *et al*., 2017; Voellmy *et al*., 2014a).

A reduced ability to perceive potential threats can also result in direct stress (Pitcher and Turner, 1986; Sutherland *et al*., 2008; Ohata *et al*., 2014). Stress occurs when individuals are unable to maintain normal physiological functions due to higher demands on the body from the environment (Schulte, 2014), in contrast to masking and distraction which affect behaviour and the efficacy of sensory system perception. Stress can be caused by temperature changes which alter the energetic states of fish, affecting activity and limiting the energy individuals have available or altering internal states so individuals in a group differ (Fisher *et al*., 2021). Stress can be measured by cortisol levels (Wysocki *et al*., 2006). Temperature changes can therefore affect the hydrodynamic benefits of shoaling, where higher temperatures result in less cohesive groups (Bartolini *et al*., 2015; Weetman *et al*., 1998). The impact of temperature is often confounded with oxygen concentrations, where in hypoxic conditions reduced shoaling is caused by a trade-off between maintaining close, cohesive shoals and accessing oxygen (Domenici *et al*., 2002; Israeli, 1996; Moss and MacFarland, 1970). Hypoxia can also be a result of eutrophication (Hagy *et al*., 2004; Rydberg *et al*., 1990), and similar impacts can be found as a response to chemical pollutants, where chemicals interfere with physiology and can reduce social interactions (Webber and Haines, 2003; Brodin *et al*., 2013; Michelangeli *et al*., 2022). The masking, distraction, and stress effects caused by environmental change rarely occur in isolation, they can influence social behaviour in different and combined ways, and field studies can help elucidate the impacts of such change on natural freshwater populations and environments.

Taking the restrictions of current techniques into consideration, and the threats faced by freshwater fish communities, we developed a method using passive funnel traps allowing us to compare fish social behaviour to environmental parameters. Here, we can quantify the aggregation and activity of fish populations based on the numbers of fish caught in each trap. Due to potential metabolic limitations we would expect to catch fewer fish at lower temperatures, but that they would also be more aggregated (i.e. less distributed across traps) (Bartolini *et al*., 2015), while being less aggregated in low oxygen conditions (Domenici *et al*., 2002). Visual constraints could also lead to fish being less aggregated when turbidity is higher (Ohata *et al*., 2014; Chamberlain and Ioannou, 2019) and where there is lower light intensity (Pitcher and Turner, 1986), due to vision being an important sensory modality for shoal cohesion (Ioannou *et al*., 2011).

Additionally, we measured the body lengths of each caught fish with the aim to quantify the phenotypic assortment of groups. Assortment occurs when there is non-random grouping of individuals that share similar characteristics, including size, and experimental studies have demonstrated a preference for fish to choose to group with those similar to them (Ranta and Lindström, 1990; Krause e*t al*., 1996; Peuhkuri *et al*., 1997; Ward and Krause, 2001). Assorting preferentially with those similar to you is potentially adaptive through reduced foraging competition (Peuhkuri, 1997) and improved predator avoidance through predator confusion (Krakauer, 1995; Ioannou *et al*., 2008).

We used three-spine sticklebacks (*Gasterosteus aculeatus*) as our model species due to their prevalent use in behavioural and genetic research, and because they are commonly found in fresh and brackish water in the UK. Sticklebacks are facultatively social and have been found to prefer to join groups of individuals similar to them in size (Ranta and Lindström, 1990). They use visual cues in their social behaviours (Huntingford and Ruiz-Gomez, 2009) and previous research has shown this can be affected by environmental conditions such as light and turbidity (Candolin *et al*., 2007; Wong *et al*., 2007).

## Materials and Methods

### 1 Study sites

The study was carried out in four ponds in Bristol, UK (Table 1). Sites were chosen based on having populations of three-spined sticklebacks (*Gasterosteus aculeatus*) adequately large to catch enough fish per site visit for analysis of aggregation and assortment by body size. Site choice was also dependant on accessibility and appropriate water depth for traps to be deployed. Sampling carried out at these sites was approved by the Environment Agency UK.

**Table 1.**
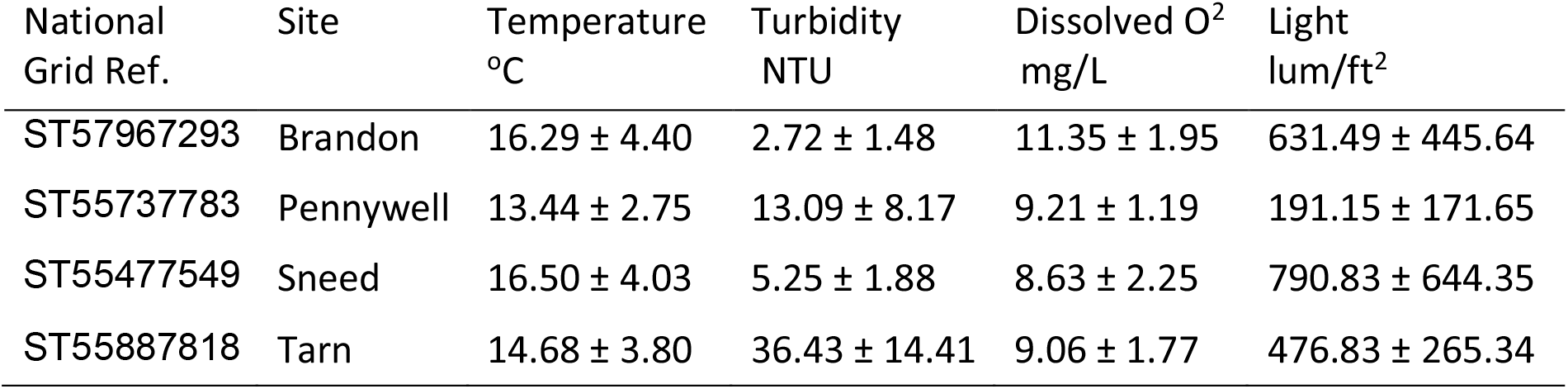
Environmental variables at each site Mean ± standard deviation of environmental variables recorded at each site. These are calculated from the average of all recordings per site visit over the course of data collection from May to November 2021.

### 2 Experimental Procedure

The study was designed to quantify the aggregation of fish populations by comparing the numbers of fish caught across five funnel traps simultaneously deployed at each visit to a site (Figure 1). Traps were deployed at equidistant locations along a pond, as much as access and water depth allowed (Figure 2); these trap locations were maintained throughout data collection and labelled 1 – 5 from left to right (see pins on Figure 2). Traps were either dropped slowly into the water using the long string attached to the trap (Figure 1) where it was possible to get close enough to the trap location, or thrown by holding two corners of the trap when the trap needed to reach a further distance, for instances where water was too shallow near the bank or there was poor access to the edge of the water. Deployment required careful positioning to ensure the temperature and light intensity logger (HOBO MX2202) attached to each trap was facing upwards in the water, and misalignment would require pulling the trap back in with the string and redeploying in the same location. Traps were not baited to avoid attracting unwanted species; traps were baited in the first week with bread, but tended to attract other species such as carp. After the two-hour sampling period where traps were left in the water undisturbed, they were pulled in using the string attached to each trap. If the trap contained fish it was quickly moved to an area of shallow water in the pond so that all caught fish were fully submerged in pond water but shallow enough to prevent escape through the openings in the trap, allowing time for counting and measuring the fish while preventing unnecessary stress.

**Figure 1.**
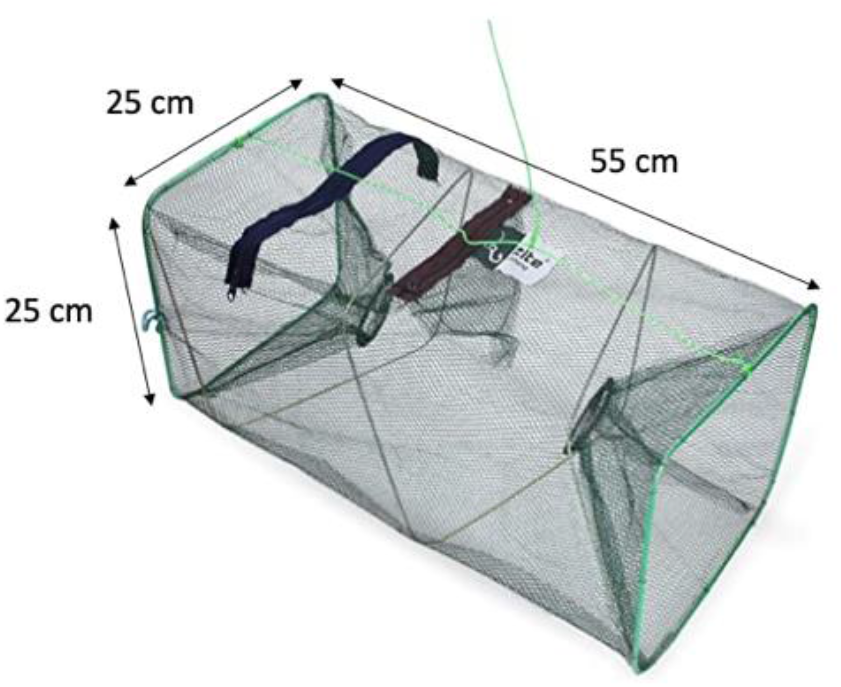
Zite fishing bait fish trap with 5mm small mesh and 2mm thick wire frame.

**Figure 2.**
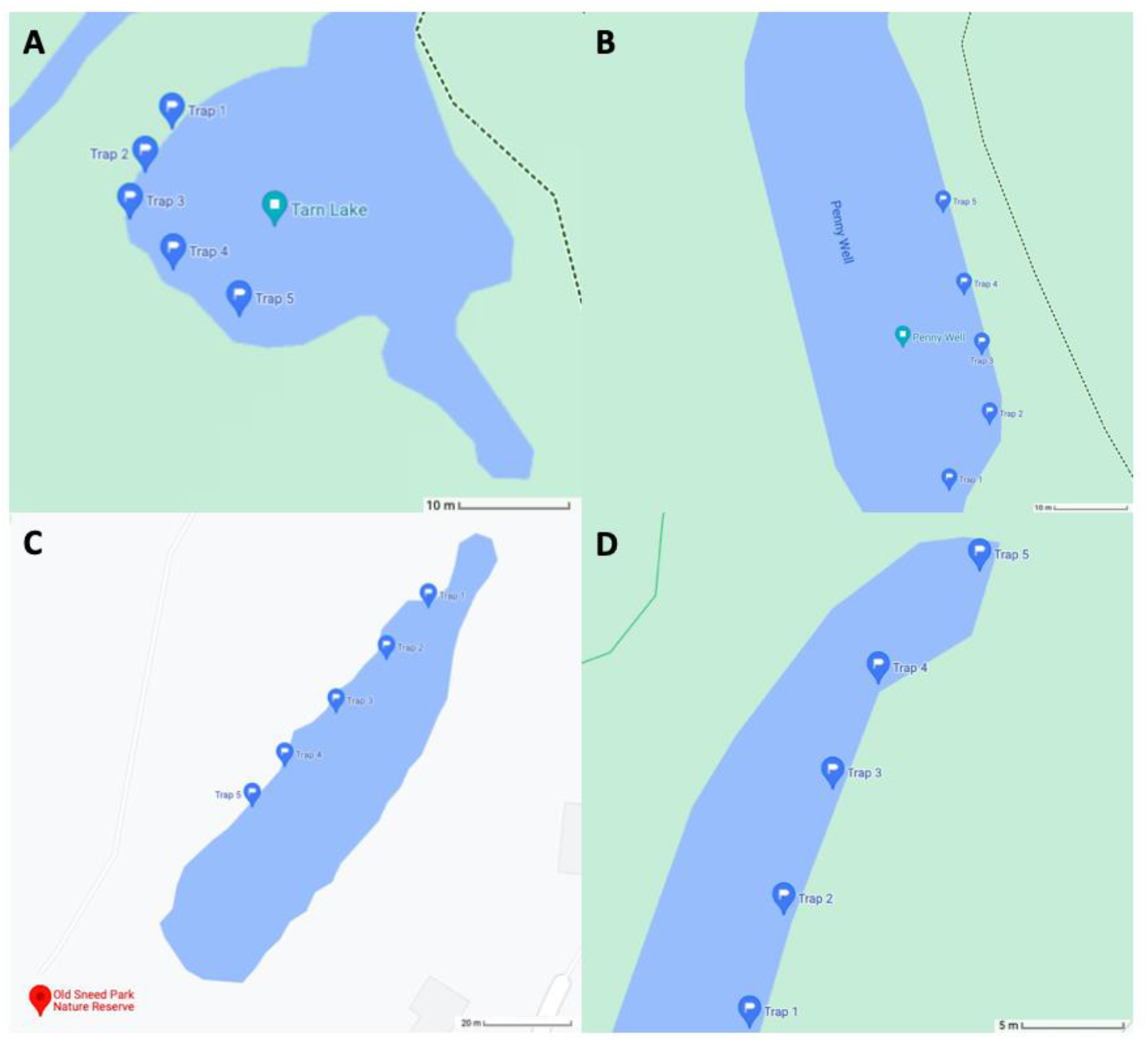
Map of each field site. Blue pins indicate trap locations from 1 – 5. **A**. Tarn: mean distance between traps 4.7 m, total size ∼0.075 hectares; **B**. Pennywell: mean distance between traps 6.7 m, total size ∼0.16 hectares; **C**. Sneed: mean distance between traps 8.8 m, total size ∼0.25 hectares; **D**. Brandon: mean distance between traps 4.9 m, total size ∼0.016 hectares. Images taken from Google Maps.

Sampling took place on alternating weeks between May and November 2021. Which site was visited on which day during a week of data collection was decided by random shuffling of the sites in R version 3.3.3 (R Core Team, 2017). Traps were left for two hours between either 10am – 12pm (AM) or 1pm – 3pm (PM) to account for any confounding effect of time of day. The first sampling time for each site was chosen by coin-flip, and alternated between AM and PM thereafter.

#### 2.1 Environmental Variables

Water temperature and light intensity were recorded using a HOBO MX2202 attached to each trap, which logged data every minute for the two-hour sampling period. Turbidity and dissolved oxygen were measured from water samples taken at each trap location, with samples taken before and after the traps were deployed (i.e. at the start and end of each site visit). Turbidity was measured using a Thermo Scientific Orion AQUAfast AQ3010 Turbidity Meter, and dissolved oxygen using a Lutron Dissolved Oxygen Meter PDO-519.

#### 2.2 Measuring Fish

Fish caught in each trap were counted, also noting the number of males in breeding condition with characteristic red colouration (Tinbergen, 1952). Body length measurements were taken by placing groups of fish in a bucket (Figure 3) with a 10cm scale bar and water from the pond to a depth of 5 cm. The number of fish in the bucket varied between 1 and 21, depending on the number of fish caught in the traps and their size (a greater number of juveniles could be imaged accurately). A GoPro Hero5 was attached to the side of the bucket, 27 cm above the water surface and oriented downward to give an overhead view of the fish. 4000 × 3000 pixel resolution images were taken in burst mode, where 10 photos were taken over two seconds, repeated multiple times if fish were closely aggregated, with the aim to capture an image where all fish are clearly visible and not overlapping. Body length measurements were then made using ImageJ (version 1.53; Schneider *et al*., 2012). This allowed for reduced stress to the fish and more efficient data collection than manual handling and measuring each fish individually, for example using callipers.

**Figure 3.**
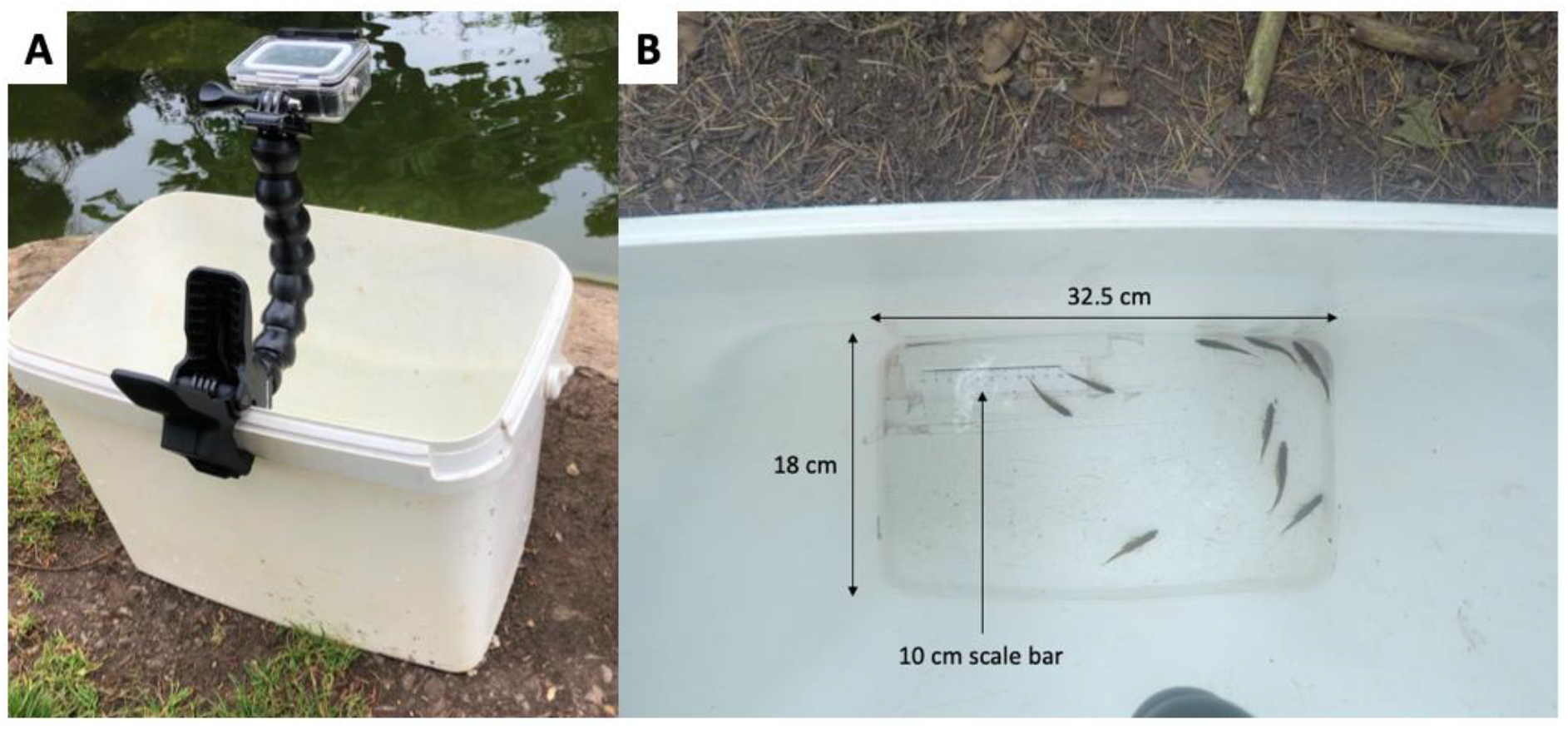
(A) Picture of bucket and camera set up; (B) example of picture with fish in bucket, subsequently used to measure body length in ImageJ (version 1.53).

#### 3 Statistical analysis

All analyses were conducted in R (version 4.1.2; Core Team, 2017) using RStudio (version 2021.9.1.375; RStudio Team, 2020). The mean of the temperatures and light intensities recorded from the start to the end of the sampling period at each trap location at one-minute intervals was calculated to give an average temperature and light intensity value for each trap location within each site for every site visit. Turbidity and dissolved oxygen measurements from the start and end of each site visit, from each trap location, were also averaged (mean). When analysing response variables which only had a single value per site visit, rather than at each trap location, for example the total number of fish caught in that visit, these variables were then averaged (mean) across the five trap locations. To test for correlations between the continuous explanatory variables (temperature, light intensity, dissolved oxygen, turbidity, and week of data collection), for both per-trap location and per-site visit values separately, relationships were tested with Spearman’s rank correlation coefficient (Spearman’s *r*_s_) in R (R Core Team, 2017).

To determine which variables could explain the variation in the total number of fish caught at each site visit, we compared 7 generalised linear mixed-effects models (GLMMs) with a negative binomial distribution and default log link function. The response variable was the total number of fish caught per visit to a site, i.e. the sum of the number of fish caught per trap, and the pond site was included as the random effect. Each of the 7 models had a different fixed effect: temperature, light intensity, dissolved oxygen concentration, turbidity, the week of data collection (1 to 13), and whether sampling occurred in the morning or afternoon (AM or PM); a null model with no fixed effect was also included in the model comparison set. We then compared the Akaike information criterion values corrected for small samples sizes (AICc) for each model using the ICtab function in R (*bbmle* version 1.0.24; Bolker and R Development Core Team, 2020). Models with lower AICc values are more likely given the data, and the model with an ΔAICc of zero is the most likely model. Explanatory variables included in models that had AICc values of greater than two units less than the null model were considered to be important predictors of the response variable, i.e. these models were considered to have strong support (Burnham and Anderson, 2002). In addition to comparison to the null model, we could also determine which explanatory variables were more likely to predict the variation in the response variable than others.

To explore these trends further, we repeated this process for the number of fish caught per trap, rather than per site visit, as a function of the same explanatory variables with the addition of trap location nested within pond site as the random effect. This allowed us to consider the environmental variation at a smaller spatial scale, at the level of the trap location. The aggregation of fish between the traps was determined by calculating the index of dispersion (i.e. the variance ÷ mean) from the number of fish caught in each trap per site visit. An aggregation score of 0 indicates fish are evenly distributed across the traps and therefore not aggregating, a value of 1 indicates fish are randomly distributed between the traps, and higher values indicate fish are more aggregated (Figure 4A). Cases were removed from the analysis when the total number of fish caught in that visit was too low to show aggregation by this measure (a threshold of 26 fish was determined from plotting the aggregation score as a function of the total number of fish caught; Figure 4A). To determine which variables were likely to predict the aggregation of fish, we compared 8 negative binomial GLMMs. The response variable was the aggregation score for each site visit, and each model had a different explanatory variable: temperature, light intensity, dissolved oxygen concentration, turbidity, week of data collection and morning or afternoon. Here, an additional model was considered, which had the proportion of red-bellied breeding condition males caught (i.e. number of red males ÷ total fish caught) as an explanatory variable. The eighth model was the null model that lacked an explanatory variable. Pond site was included as the random effect in all models. AICc values were then compared in the same way as previously described.

**Figure 4.**
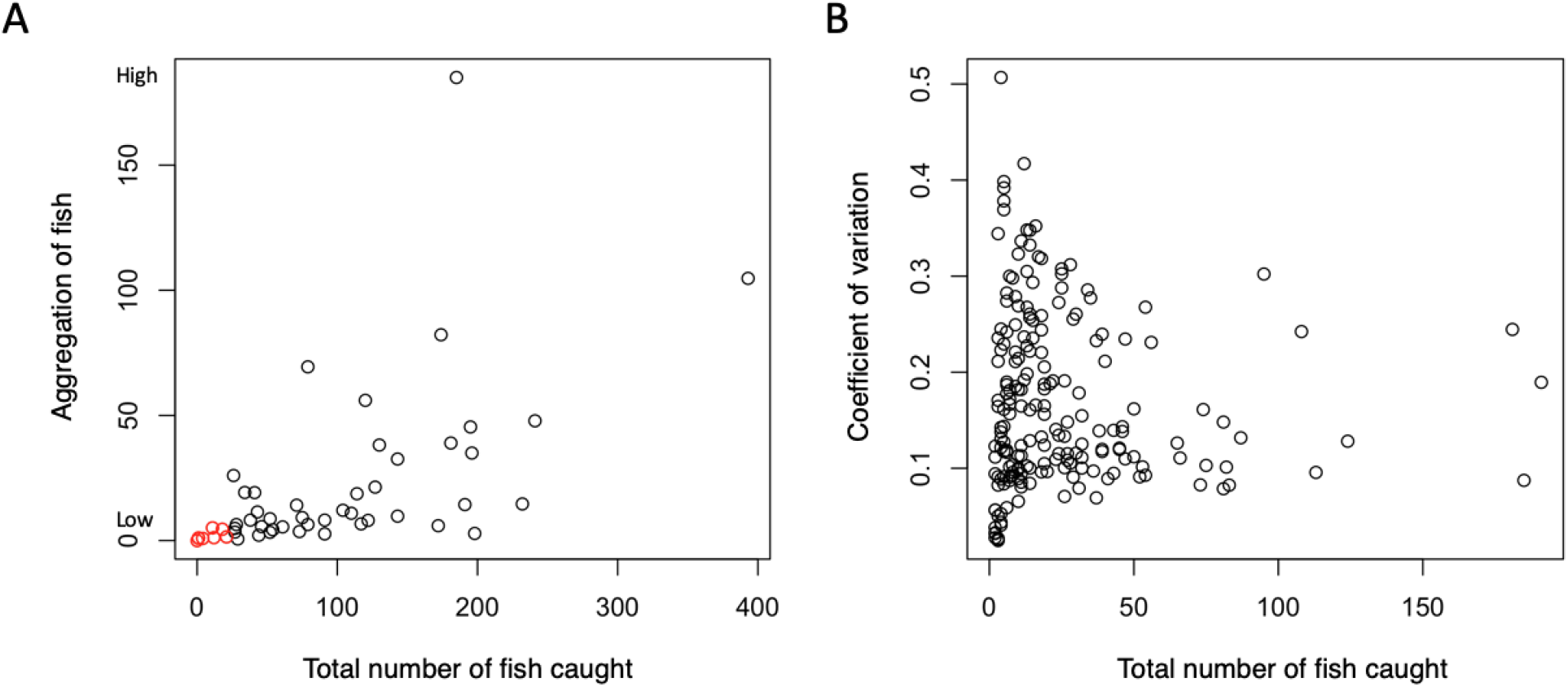
(A) Aggregation scores (index of dispersion, variance ÷ mean) for each site visit as a function of the total number of fish caught at that site visit. Larger values on the y axis represent cases where fish are more aggregated. Red points indicate data that was removed from the analysis, as the aggregation score did not seem to be able to take large values when the total number of fish caught was <26. (B) Coefficient of variation (COV) values for each trap compared to the number of fish caught in that trap. Larger values on the y axis indicate high variation in body length among fish in a trap.

All GLMMs were run using the glmmTMB function (*glmmTMB*; Brooks *et al*., 2017). The assumptions of all models were verified using QQ plots and dispersion tests using the residual diagnostics for mixed regression models (DHARMa; Hartig 2019).

As a measure of phenotypic assortment, we used the coefficient of variation (COV; i.e. standard deviation ÷ mean) of body length, calculated for the fish caught in each trap, excluding cases where only one or no fish were caught in the trap (Figure 4B; Croft *et* al., 2009). A COV of zero would indicate there is no variation in size between the fish in a trap; the higher a COV value, the more variation (relative to the mean body size) there is in body size between fish in a trap. The median COV of all the traps at one site visit was used as the observed COV for each site visit (observed siteCOV). To determine whether fish caught in the traps were more or less phenotypically assorted by body length than expected by chance, we calculated the expected median COV for each site visit, assuming random assortment within a visit to a site. The expected value for assortment was calculated using a constrained randomisation procedure in which individuals caught across traps on one site visit were randomly re-distributed across the five traps, maintaining the number of fish caught per trap as in the observed data. The COV of body length was calculated per trap from each resampling, and the median of these values were saved as the value of expected assortment (expected siteCOV). This was iterated 10,000 times for each site visit and an expected distribution of assortment values was generated. The observed siteCOV was used as a quantile on the corresponding expected distribution.

In the two-tailed tests, quantiles are statistically significant when < 0.025, indicating positive assortment (i.e. fish are assorting with those that are similar to themselves), or when > 0.975, indicating negative assortment (i.e. fish are assorting to those that are different to themselves).

## Results

### Correlation between explanatory variables

Correlations between the continuous explanatory variables included in the models showed evidence of correlation between temperature and light intensity (Figure 5A; Spearman’s rank correlation coefficient: *r*_*s*_ = 0.60, *p* < 0.0001, n = 52), week and light intensity (*r*_*s*_ = -0.60, *p* < 0.0001, n = 52), and week and temperature (*r*_*s*_ = -0.58, *p* < 0.0001, n = 52). Correlation coefficients were similar when using the values measured at each trap location per visit (Figure 5B).

**Figure 5.**
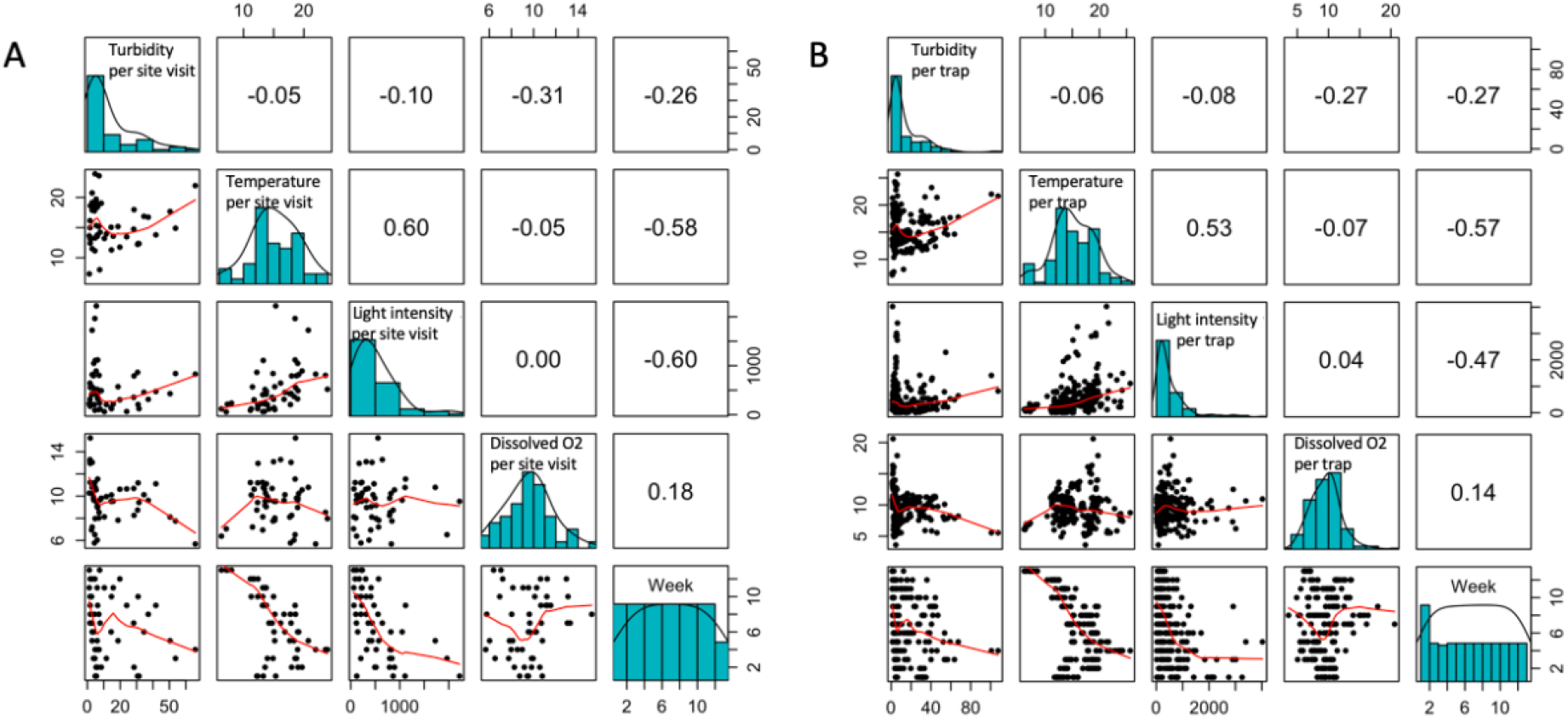
Distributions of, and correlations between, the 5 continuous variables used as explanatory variables in the models. Correlation coefficients are Spearman’s *r*_*s*_. The red curve in the scatter plots are LOWESS smoothed curves. In (A), data is averaged across trap locations giving a value for each site visit. In (B), data includes values of each variable for each trap location per site visit, thus considers within-site variation.

### Total fish caught per site visit

Analysis of the total number of fish caught at each site visit showed that the model with week as the explanatory variable was the most likely given the data (Table 2). The total number of fish caught decreased over consecutive weeks of data collection (Figure 6A). The model with temperature as the explanatory variable was less well supported, but being greater than 2 AICc units less than the null model still provides strong evidence that temperature was having an effect on the number of fish caught (Table 2). In this case, the number of fish caught increased in warmer temperatures (Figure 6B). There was some support for the model with light intensity as the explanatory variable, which was 1.1 AICc units less than the null model. Conversely, models where the only explanatory variable was dissolved oxygen, turbidity or the time of day were not supported, indicating that the number of fish caught on a site visit was not associated with variation in these parameters.

**Table 2.**
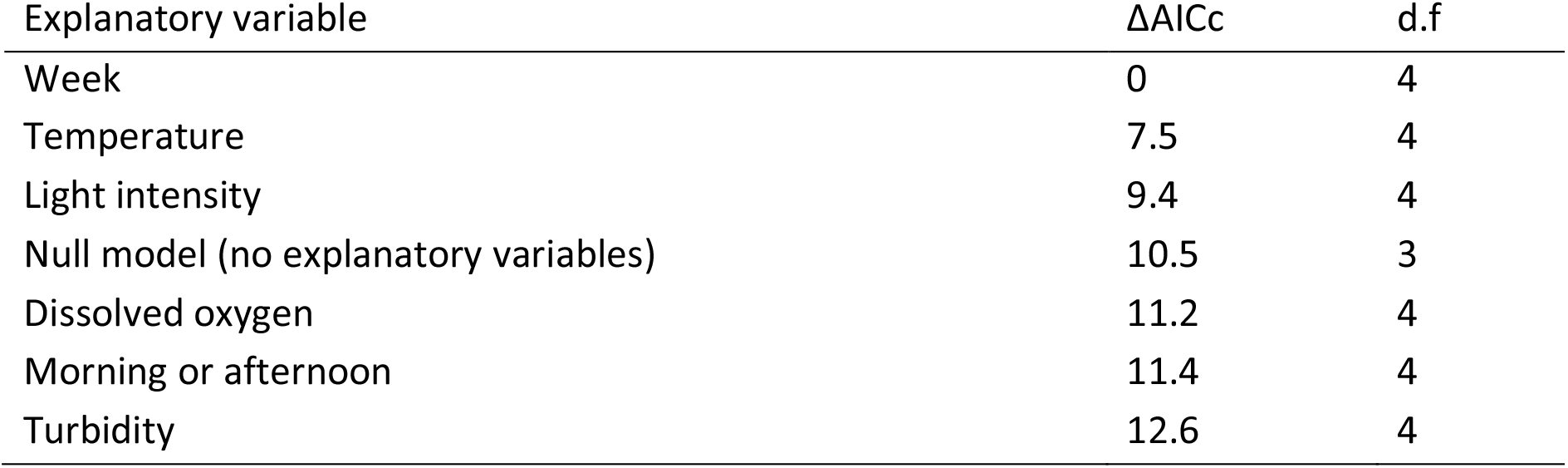
The ΔAICc for models explaining variation in the total number of fish caught per site visit. Models differ in the explanatory variable included, and all include pond site as the random effect. The null model has no explanatory variable, only the random effect. Week (week 1-13) and morning or afternoon (sampling in AM or PM) represent when data collection occurred.

| Explanatory variable | $\Delta AICc$ | d.f |
| --- | --- | --- |
| Week | 0 | 4 |
| Temperature | 7.5 | 4 |
| Light intensity | 9.4 | 4 |
| Null model (no explanatory variables) | 10.5 | 3 |
| Dissolved oxygen | 11.2 | 4 |
| Morning or afternoon | 11.4 | 4 |
| Turbidity | 12.6 | 4 |

**Figure 6.**
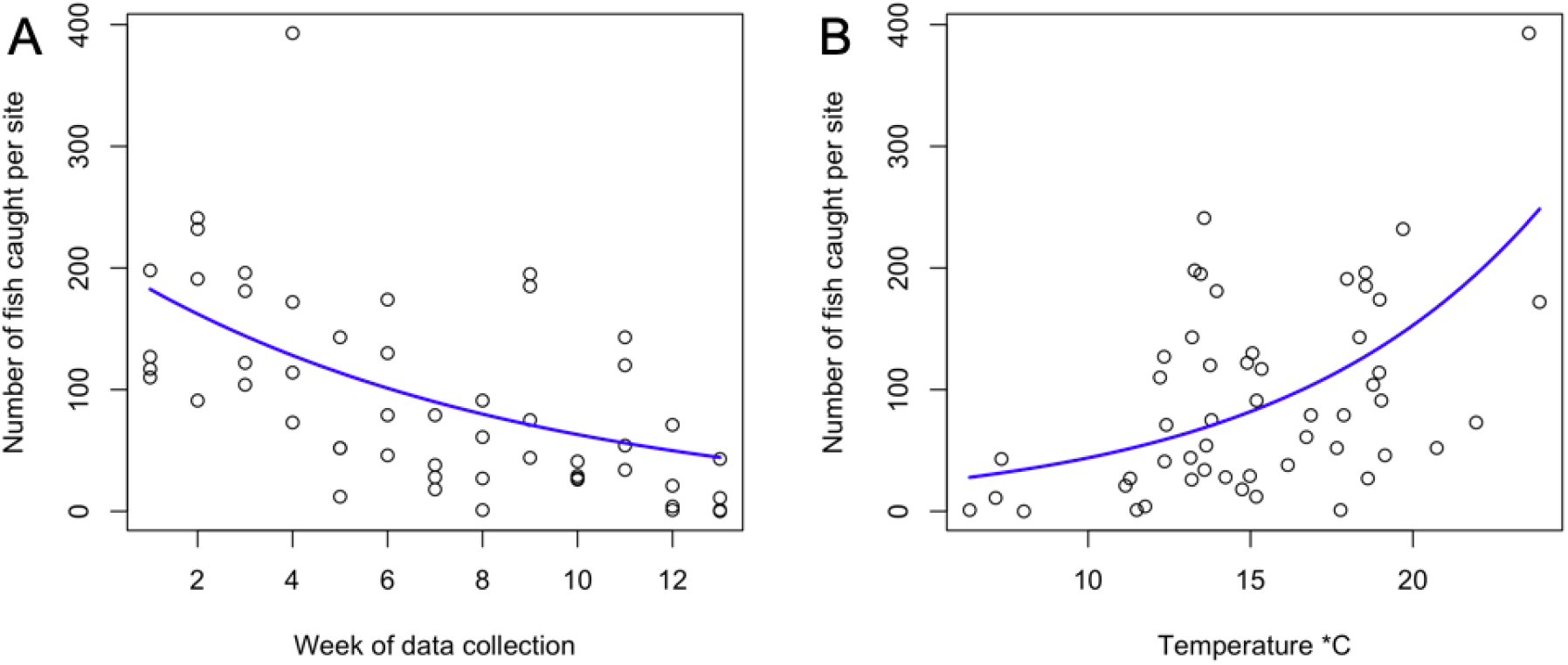
The number of fish caught per site visit as a function of (A) the week of data collection and (B) the average water temperature across the trap locations. Plotted lines show the fitted relationships from GLMM coefficients.

### Total fish caught per trap

Analysis of the total number of fish caught per trap, which takes into account environmental variation at each trap location, revealed similar trends to when only averages across a site on a visit were considered. Week was again the most likely model given the data (Table 3), where the number of fish caught decreased over consecutive weeks of data collection (Figure 7A). The model with temperature was also supported by the data, with the trend again showing the number of fish caught increased with warmer temperatures (Figure 7B). Based on the AICc being higher than the null model, there was no evidence for turbidity or time of data collection having an effect on the number of fish caught at a trap location.

**Table 3.** The ΔAICc for models explaining the total number of fish caught per trap. Models differ in the explanatory variable included, and all include trap location nested within pond site as the random effect. The null model has no explanatory variable, only the random effects. Models with environmental variables include those variables recorded at each trap location. Week (week 1-13) and morning or afternoon (sampling in AM or PM) represent when data collection occurred.

| Explanatory variable | $\Delta AICc$ | d.f |
| --- | --- | --- |
| Week | 0 | 5 |
| Dissolved oxygen | 18.5 | 5 |
| Temperature | 27.5 | 5 |
| Light intensity | 31.3 | 5 |
| Null model (no explanatory variables) | 35.6 | 4 |
| Morning or afternoon | 36.2 | 5 |
| Turbidity | 37.6 | 5 |

**Figure 7.**
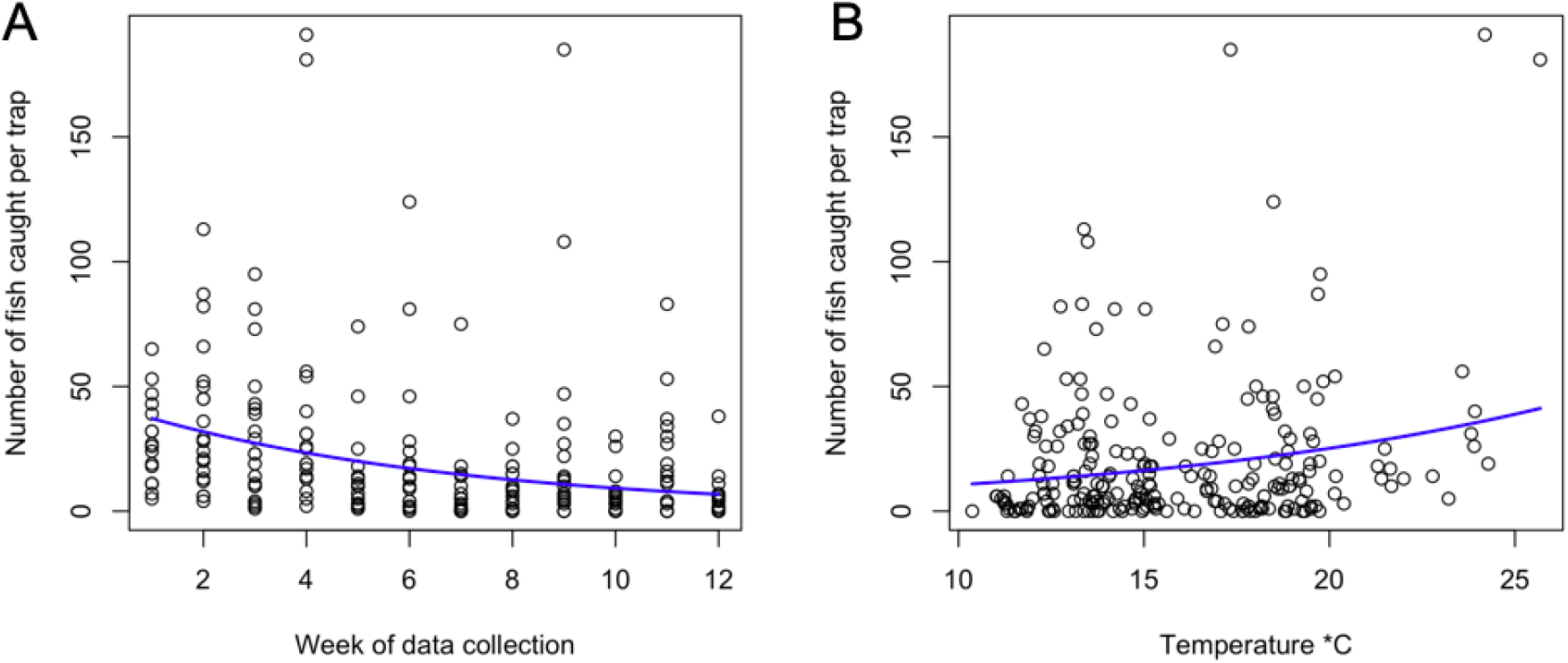
The number of fish caught per trap as a function of (A) the week of data collection and (B) the water temperature at each trap location. Plotted lines show the fitted relationships from GLMM coefficients.

In contrast to the analysis based on the averages from each site visit, the model with dissolved oxygen as the explanatory variable was strongly supported compared to the null model (Table 3). More fish were caught when the concentration of dissolved oxygen in the water was lower (Figure 8A). Light intensity was less well supported than models with some of the other variables, but with the AICc being greater than 2 less than the null, there was still strong evidence of it being likely to have an effect. Here, the number of fish caught in a trap increased as light intensity increased (Figure 8B).

**Figure 8.**
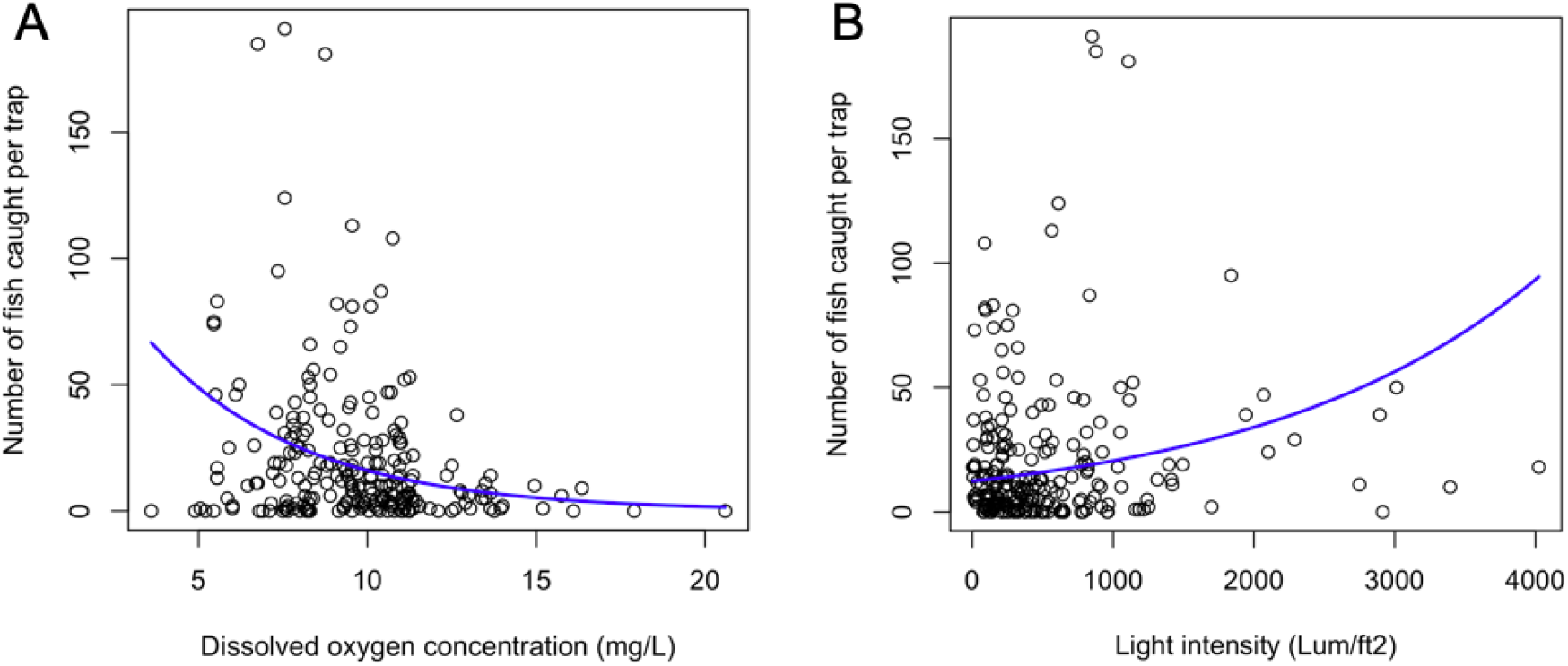
The number of fish caught per trap as a function of (A) the concentration of dissolved oxygen and (B) the light intensity at each trap location. Plotted lines show the fitted relationships from GLMM predicted values.

### Aggregation of fish

When testing the variables that predict the aggregation of fish across the traps at each site visit, the most likely model given the data was the model with the proportion of red-bellied males as the explanatory variable (Table 4). The proportion breeding-condition males was negatively associated with the aggregation score (Figure 9). All other models had larger AICc values than the null model so were not supported by the data.

**Table 4.** The ΔAICc for models explaining the aggregation of fish. Models differ in the explanatory variable included, and all include pond site as the random effect. The null model has no explanatory variable, only the random effect. Week (week 1-13) and morning or afternoon (sampling in AM or PM) represent when data collection occurred. Proportion of red males caught represents how many breeding-condition males were caught relative to the total number caught.

| Explanatory variable | $\Delta AICc$ | d.f |
| --- | --- | --- |
| Proportion of red males caught | 0 | 4 |
| Null model (no explanatory variables) | 2.6 | 3 |
| Dissolved oxygen | 3.3 | 4 |
| Morning or afternoon | 3.5 | 4 |
| Turbidity | 4.0 | 4 |
| Temperature | 4.9 | 4 |
| Week | 4.9 | 4 |
| Light intensity | 5.0 | 4 |

**Figure 9.**
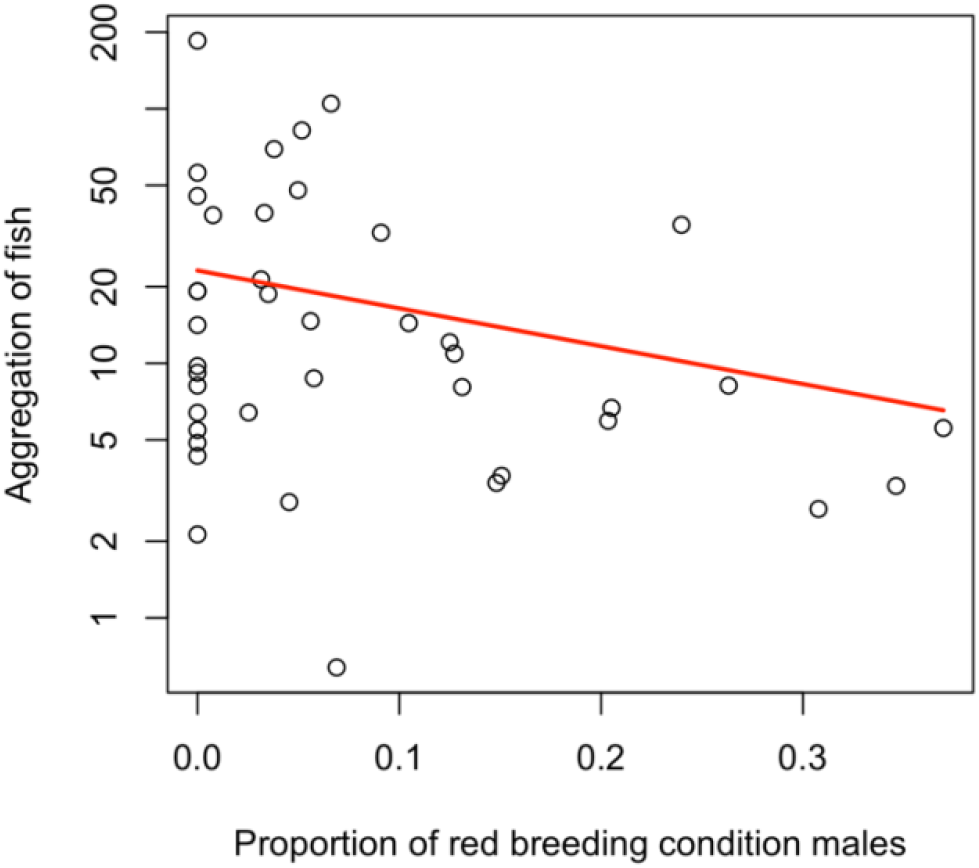
Relationship between the proportion of red-bellied males caught and the aggregation of fish. Plotted line shows the fitted relationship from GLMM coefficients.

### Body size assortment of shoaling fish

We found little evidence of body length assortment among the fish caught in the traps. Out of 44 quantiles, calculated for each site visit (where 8 visits were unsuitable for analysis due to low numbers of fish caught during the visit or the distributions of fish across traps), only 1 quantile was below <0.025 (Figure 10A). The majority of visits yielded non-significant tendencies to be positively assorted (e.g. Figure 10B); in 8 of the 44 visits, the values tended towards negative assortment (e.g. Figure 10C) but none were statistically significant.

**Figure 10.**
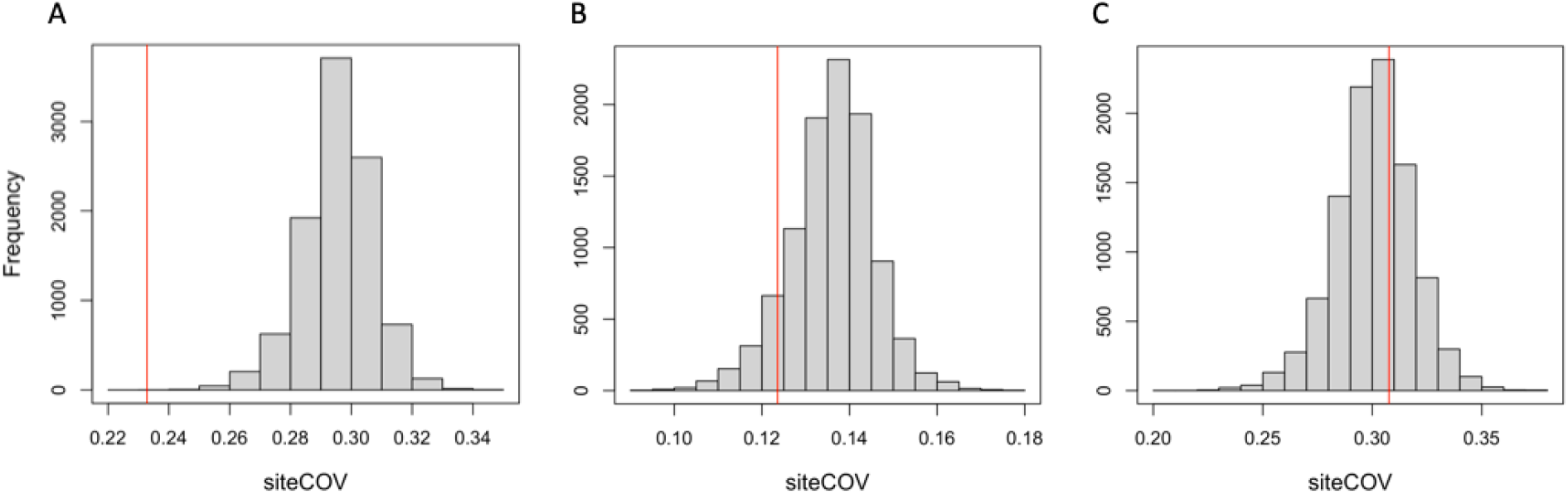
Example expected distributions generated by randomised sampling for three site visits. Red line shows observed value, i.e. the median of the COV for that site visit. Examples of (A) the statistically significant value for positive assortment (quantile 0.004), (B) a non-significant value for positive assortment (quantile 0.195) and (C) a non-significant value for negative assortment (quantile 0.68).

## Discussion

Fewer fish were caught over the weeks of data collection, likely a result of the changing seasons, where more fish were caught in spring and summer, declining through autumn and into winter. This is supported by the correlation between consecutive site visits and decreasing temperature and light intensity. Similarly, fewer fish were caught with decreasing temperatures. This can be explained by fish being ectothermic and having lower metabolisms at lower temperatures (Clarke and Johnston, 1999); therefore, in colder weather they are expected to be less active (Bartolini *et al*., 2015) and have a lower need to explore their environment looking for food (Clarke and Johnson, 1999). We found weaker evidence for reduced light intensity causing less fish to be caught. This trend could be attributed to the correlation between temperature and light intensity, where increased light intensity results in increased temperatures, and in turn increased temperature led to more fish being caught (Ilha *et al*., 2018).

The trend between the numbers of fish caught through the weeks of data collection and with decreasing temperatures was maintained when considering environmental variation at trap locations. However, this within-site variation revealed evidence for light intensity and dissolved oxygen to be impacting the numbers of fish caught. For light intensity, this could be a result of differences in canopy cover at different trap locations (canopy cover was not recorded) (Ilha *et al*., 2018). Additionally, fewer fish were caught in areas with a higher dissolved oxygen concentration. As dissolved oxygen concentration was not correlated with temperature or the week of data collection, its effect cannot be attributed to these variables. The effect only being found when considering differing oxygen concentrations within a pond site, i.e. from different trap locations, suggests fish respond to small-scale variation in oxygen concentration at the different trap locations. The effect not being found at the site level could be due to a lack of variability in dissolved oxygen concentrations between site visits. While studies have shown fish are less active at lower oxygen concentrations, the levels we have recorded are not low enough to be detrimental to the point where they would cause stress (Kramer, 1987). For example, Moss and McFarland (1970) only found measurable changes when near-lethal oxygen levels were reached or levels dropped rapidly, whereas our sites had consistent changes within normal boundaries. One possibility is that dissolved oxygen concentration is correlated with other parameters we did not record (Hagy *et al*., 2004, Rydberg *et al*., 1990).

Sticklebacks were less aggregated when the proportion of red-bellied breeding condition males caught was higher, indicating they are being less social during the breeding season. When in breeding condition male sticklebacks maintain territories, in which they build and defend nests where females are courted to lay eggs (Tinbergen, 1952). As a result of these behaviours, male sticklebacks are more solitary when in breeding condition and more aggressive towards each other, with agonistic behaviours increasing through the breeding cycle (Huntingford, 1976; Kynard, 1978). This would explain why sticklebacks were less aggregated when more of these males were present. We did not find that aggregation was related to any of the environmental variables we recorded, unlike laboratory-based studies testing a range of environmental variables (Domenici *et al*., 2002; Bartolini *et al*., 2015; Chamberlain and Ioannou, 2019; Fisher *et al*., 2021). This may be because under field conditions with variation in multiple, possibly interacting, environmental parameters, the effects of any single stressor is difficult to detect in comparison to highly-controlled laboratory studies. This may also explain why the week of data collection was also not related to aggregation, while repeatedly testing shoals of sticklebacks under laboratory conditions does show a reduction in aggregation over time (MacGregor and Ioannou, 2021).

We found little evidence for phenotypic assortment by body size in these stickleback populations. This may be a result of low predation pressure in these sites (Croft *et al*., 2009), however this is not something we measured. An additional explanation is that when populations are small, fish face a trade-off between being in a large group with many dissimilar individuals or small groups of assorted individuals. In this case, the benefits of being in a larger group could outweigh the benefit of being in an assorted one, resulting in a lack of size assortment. It is also possible that more than one shoal were caught in a trap during the 2 hours trial time, where each shoal was assorted but there was no assortment between different shoals caught in the same trap (i.e. shoals were not more likely to enter a trap already containing a shoal that had a similar, compared to a dissimilar, body size distribution). Considering this, future use of our sampling method could be adapted to focus on measuring assortment by altering the amount of time traps are left in the water, or by coupling the method with video recording when fish enter. Video recordings could be used to exclude cases where multiple shoals have entered the trap within a sampling period from the analysis of assortment, or live monitoring via video could be used to remove the traps once a shoal has entered.

A possible source of bias in our method is for certain behavioural characteristics to be over- or under-represented due to using passive traps (Wilson *et al*., 1993; Biro and Dingemanse, 2009). Kressler *et al*. (2021) found that when using passive traps, more active fish were captured sooner, and counter-intuitively, when traps contained conspecifics, less-social fish were also captured sooner. This is something to be considered, and could even provide an interesting opportunity for further work using this method, to explore personality of the sampled fish in the context of sociality and changing environments. A related issue is the novelty of traps (Wilson *et al*., 1993; Michelangeli *et al*., 2016; Kressler *et al*., 2021), but because we were resampling the same sites over months with the same traps, it is likely that the traps are no longer considered to be a novel stimulus. As fewer fish were captured over the weeks of data collection, it could be inferred that sticklebacks are learning to avoid the traps over time. Again this could be examined in future work; observations of how the fish enter traps and whether they appear to actively avoid them would be necessary to suggest that sticklebacks are learning to not enter traps.

In future applications, this method could be used to compare sites with distinct differences, for example, polluted versus pristine environments. The effects of environmental factors other than the ones we have looked at here, such as chemical pollutants (Michelangeli *et* al., 2022) or anthropogenic noise (Sarà *et al*., 2007; Purser and Radford, 2011; Voellmy *et al*., 2014a, 2014b), could also be investigated, as well as considering multiple stressors (Ormerod *et al*., 2010; Ginnaw *et al*., 2020). Additionally, hypotheses could explore more biotic factors in the environment, such as predation risk (Ioannou, 2021) or invasive species (Strayer *et al*., 2006). In the case of invasive species, the association of native and invasive species within traps could be used to infer whether these species are interacting socially or are avoiding one another (Morelia *et al*., 2014), and whether environmental parameters affect these interactions (Glotzbecker *et al*., 2015).

Here we have established a novel field method for investigating the impacts of environmental variation on social behaviour in fish. Our results demonstrate the validity of the method, presenting trends that we would expect to find in stickleback populations based on breeding condition and external abiotic factors. This method has some substantial benefits over other techniques, particularly being cost-effective and feasible practically. However, it does come with caveats relating to the size of groups, species, and other potential biases which may be revealed through further investigation. Nevertheless, it provides an opportunity to investigate hypotheses regarding fish social behaviour under environmental change, anthropogenic pollution, and other biotic factors under field conditions, questions which to date have been dominated by experimental laboratory studies (Fisher *et al*., 2021).

## Acknowledgments

This research was funded by a Natural Environment Research Council grant no. NE/P012639/1 awarded to C.C.I. We would like to thank Costanza Zanghí for assistance with fieldwork organisation and Iestyn L. Penry-Williams for advice on data analysis.

## Author contributions

M.A.C. developed the experimental design, undertook data collection and data analysis, and authored the manuscript; C.C.I. conceived the project and provided supervision of M.A.C.

